# Lever arm Flexibility Controls the extent of (Un)coupling to the Motor Domain in Myosin Motors

**DOI:** 10.1101/2025.08.07.669198

**Authors:** Mauro L. Mugnai, Yonathan Goldtzvik, D. Thirumalai

**Affiliations:** Department of Chemistry, The University of Texas at Austin, Austin, Texas, USA; Department of Structural and Molecular Biology, University College London, London, UK

## Abstract

Dimeric myosin motors, with both heads simultaneously bound to filamentous actin, are in a frustrated conformation. The lever arm of each head would prefer to orient forward, but the inter-head tension hinders the relaxation to the state favored by a myosin monomer. Here, we investigate theoretically the impact of lever arm stiffness and coupling to the head domain using a polymer model. The theory for MV and MVI qualitatively reproduce the salient experimental observations. Furthermore, we construct chimeras in which the lever arm and head domains are swapped, and predict that the fluctuations of the LH lever arm of MVI are strongly impacted by the stiffness of the lever arm. Finally, by continuously and independently varying the lever arm persistence length and the strength of its coupling to the head domain we predict their roles in altering the average geometry and conformational flexibility of the dimer. We explore conditions under which LH lever arm flexibility and angle with respect to F-actin lead to coupling to the motor domain.

For Table of Contents Use Only

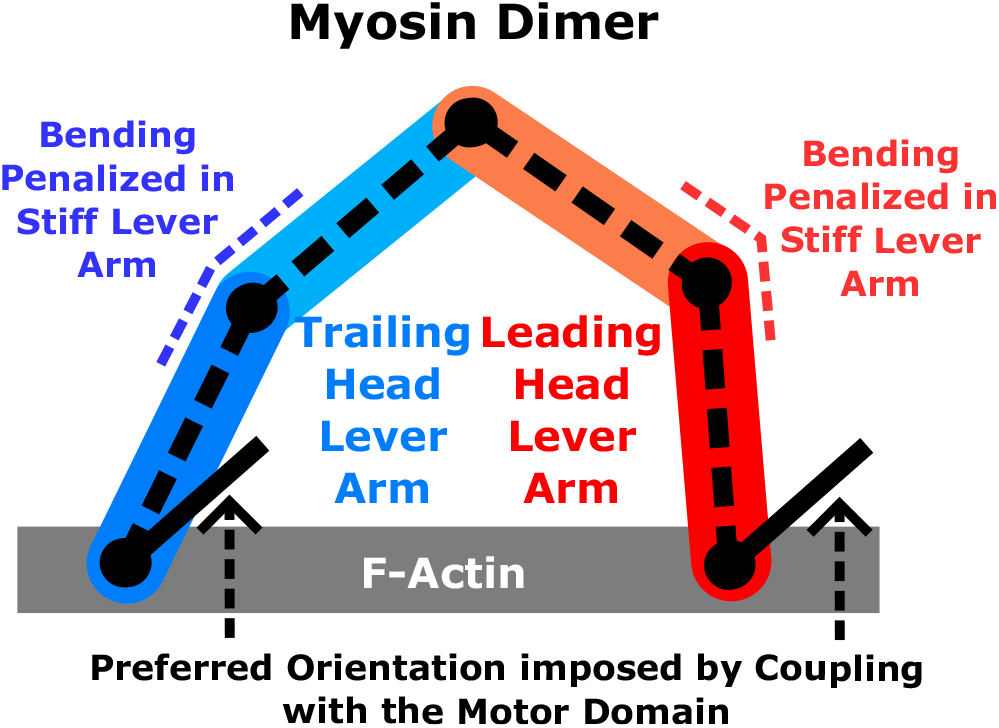

## Introduction

Dimeric processive myosin motors, such as myosin V (MV) and myosin VI (MVI) use the energy released by the hydrolysis of ATP in order to take multiple ≃36 nm step along the filamentous actin (F-actin) hand-over-hand (1–7). Both motors have a large duty ratio, which means they spend a substantial fraction of the duration of the ATPase cycles tightly bound to the actin filament (8, 9). This implies that the two-head-bound (2HB) conformation is highly populated, which makes it important to characterize the structure of the 2HB state in order to understand differences between the MV and MVI.

In a minimal description, myosin motors are modeled as a motor (or head) domain that binds F-actin and hydrolyzes ATP. During the catalytic cycle, myosins undergoes a power stroke the rotation of the motor domain results in the forward motion. The motor domain is ≃5 nm in size, and thus in order for a myosin dimer to advance by ≃36 nm per step, the rotation of the motor is extended by an oblong domain called “lever arm”. Electron microscopy (EM) (10) directly revealed that the two heads of myosin V (MV), the archetypal processive myosin motor, are bound to filamentous actin for a large fraction of the catalytic cycle. EM and subsequently atomic-force microscopy (AFM) (4) showed that the leading head (LH) of the MV dimer is under tension, as evidenced by the backward bending of the stiff lever arm, which ≈ 20 nm-long and is made of 6 IQ domains bound to calmodulins (CaMs). The tension arises due to a combination of the LH power stroke, which favors the re-orientation of the LH lever arm in the forward direction along F-actin, and the presence of the trailing head (TH), which is bound to actin at a distance, *d* = 36 nm from the LH and therefore hinders the forward swing of the LH lever arm. The combination of these two factors dictates the measured orientation of the lever arm with respect to actin (see Hinczewski *et al*. (11) for a detailed discussion).

Myosin VI (MVI) is another processive member of the myosin superfamily, which differs from MV in several aspects. First, MVI walks in the opposite direction (traverses from the + to − end along F-actin (12)) in part due to a structural unit, the ins2 domain, that acts as a reverse gear (13, 14). Second, the lever arm of MVI is made of the ins2 (bound to a CaM), of a single CaM-bound IQ domain, that is much shorter than the lever arm of MV. The existence of a lever arm extension (LAE) is therefore necessary in order to explain the average step size, which is approximately the same in both the motors. Because the lever arm is “hidden” in MVI, its nature has been debated (15– 20). One possibility is that a helical bundle called the proximal tail (PT) could serve as a LAE. According to this model, the PT would unfold upon dimerization, and attain an extended conformation possibly stabilized by binding a CaM (16, 19, 20). In an alternative scenario, the lever arm might be extended by the medial tail (MT), a single alpha helix (SAH) stabilized by internal salt bridges (15, 21, 22). Regardless of the mechanism, it is likely that MVI lever arm is more flexible than the MV counterpart (23). Third, the step-size distribution in MVI is broad, which includes frequent inchworm-like steps and backward steps (5). In contrast, MV rarely takes a backward step in the absence of resisting load (24). Fourth, a number of experiments have identified that the lever arm of the LH in MVI displays a considerable amount of conformational flexibility, which was interpreted as pliancy leading to the uncoupling from the motor domain during the stepping process (2, 14, 25–27).

The differences between MV and MVI sketched above behooves us to understand the relation between the lever arm flexibility and the coupling to the motor domain, which has implications on stepping kinetics of the two motors. Because of tension between the two heads in MV, which results in a backward bending of the LH lever arm seen in experiments (4), the lever arm movement is coupled to the motor motor domain. Thus, the fluctuations in the conformations of the lever arm are restricted in the 2HB state. In contrast,following an important study (27), we used coarse-grained Molecular Dynamics (MD) simulations to show that the uncoupling of LH from the motor domain in MVI is a consequence of a two-step mechanism for the power stroke (28). In the first step, the motor domain attains the post-stroke conformation, while the lever arm points backward. In the second step, the lever arm completes the power stroke (28). The presence of the actin-bound TH would prevent the second step until detachment of the TH, resulting in the LH lever arm undergoing large fluctuations, as noted in experiments (2, 25).

Our goal is to evaluate how (i) lever arm flexibility and (ii) uncoupling from the motor domain contribute to the conformational freedom of the LH lever arm in MV and MVI. We developed a polymer model of a myosin dimer with both heads bound to actin. Our model is similar to the dimeric constructs for myosin V (11, 29, 30) and myosin VI (31). Using the structural information from a number of experiments (see caption to Table 1), we determined the parameters of the model and compared the range of conformations accessible to the lever arms of MV and MVI. We also analyzed two chimeras in order “continuously” change the stiffness the of the lever arm and the strength of the lever arm coupling to the motor domain. We show that the average angle of the LH lever arm, with respect to actin, is increased by the coupling to the motor domain, and decreased as the lever arm becomes stiffer. In contrast, the increased coupling and stiffness reduce the extent of fluctuations the LH lever arm. Our theory aids the interpretation of the 2HB conformation extracted from experiments, because it can be used to understand whether the structures imply lever arm flexibility and strength of coupling between lever arm and motor domain.

**Table 1:** Parameters in the model. The third column refers to MV, the fourth to MVI, the fifth to a chimera in which the motor domain is made of MV and the lever arm of MVI, and the last column describes to a construct in which the motor domain is taken from MVI and the lever arm from MV. **MV:** The contour length was estimated from the distance between the alpha carbons of residues 755 and 909 in the structure of the inactivated myosin V (PDBID: 2DFS, from Liu *et al*. (35)), which covers the whole lever arm (from the end of the converter to the 6 IQ domains and related CaMs). The value is close to the 26 nm reported by Sun and Goldman (23). It is in excellent agreement with the 31 nm listed by Walker *et al*. (10) if one includes ≈ 7.5 nm (distance from residue 373 and residue 755 in PDB 2DFS) for the motor head. The persistence length is taken from Hinczewski *et al*. (11). Although the value suggested elsewhere (23) is lower (≈150 nm), in both cases *L*/ *l*_*p*_>> 1, so it is not expected to make a significant difference. The TH lever arm angle with respect to F-actin is taken from the average obtained by Kodera *et al*. (4) for the single-head images of MV in the presence of 1 mM ADP. The value is not far from the 40° reported by Walker *et al*. (10) for the TH. The LH angular constraint is chosen so that the angular distribution of the LH is peaked around 115°, the value reported by Walker *et al*. (10). The strength of the constraint, 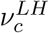, (in *k*_*B*_*T* units is less than the value used previously (11). Note that here we use the OBR model for the lever arm, which is less accurate than the WLC adopted in (11). In the OBR model, the constraint of the head impacts half of the length of the lever arm, which is much more than its effect in WLC. This might account for the discrepancy between the two. **MVI:** The length of the lever arm is estimated as a combination of the length of ins2 and the IQ domain [≈7.5 nm (28)], the length of the PT [≈3 nm (15)], and a fully-helical and straight (22) medial tail of ≈10 nm [66 residues (15) times 0.15 nm per residue]. The combination of these three units is shown in Fig. A2b. The persistence length of MVI is taken as the value obtained for the MT (22), which is likely smaller than the persistence length of the CaM-bound ins2 and IQ domain and the folded PT. The lever arm angle is estimated from the cryo-EM image of an ADP- and actin-bound myosin provided by Gurel *et al*. (36) (PDBID: 6BNW). The angle in the table is between the portion of ins2 resolved (residues 774-797) and the actin filament. The LH is assumed to be “uncoupled” from the motor domain, and therefore the angular constraint 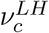 (in unit of *k*_*B*_*T*) is taken to be 0.

| parameter | symbol | MV | MVI | MVmd/MVlla | MVImd/MVla |
| --- | --- | --- | --- | --- | --- |
| Contour Length | $L$ | 23.0 nm | 21.0 nm | 21.0 nm | 23.0 nm |
| Persistence Length | $l_p$ | 310 nm | 22.4 nm | 22.4 nm | 310 nm |
| TH angle with F-actin | $\theta_T$ | 31° | 25° | 31° | 25° |
| LH angular constraint strength | $\nu_c^{LH}$ | 50 | 0 | 50 | 0 |

## Results

### Once-Broken Rod Model

We model the lever arm of MV and MVI as two rigid rods connected by a restrained hinge (Once-Broken Rod, OBR, see Fig. 1a). The energy of such a conformation is,

**Figure 1.**
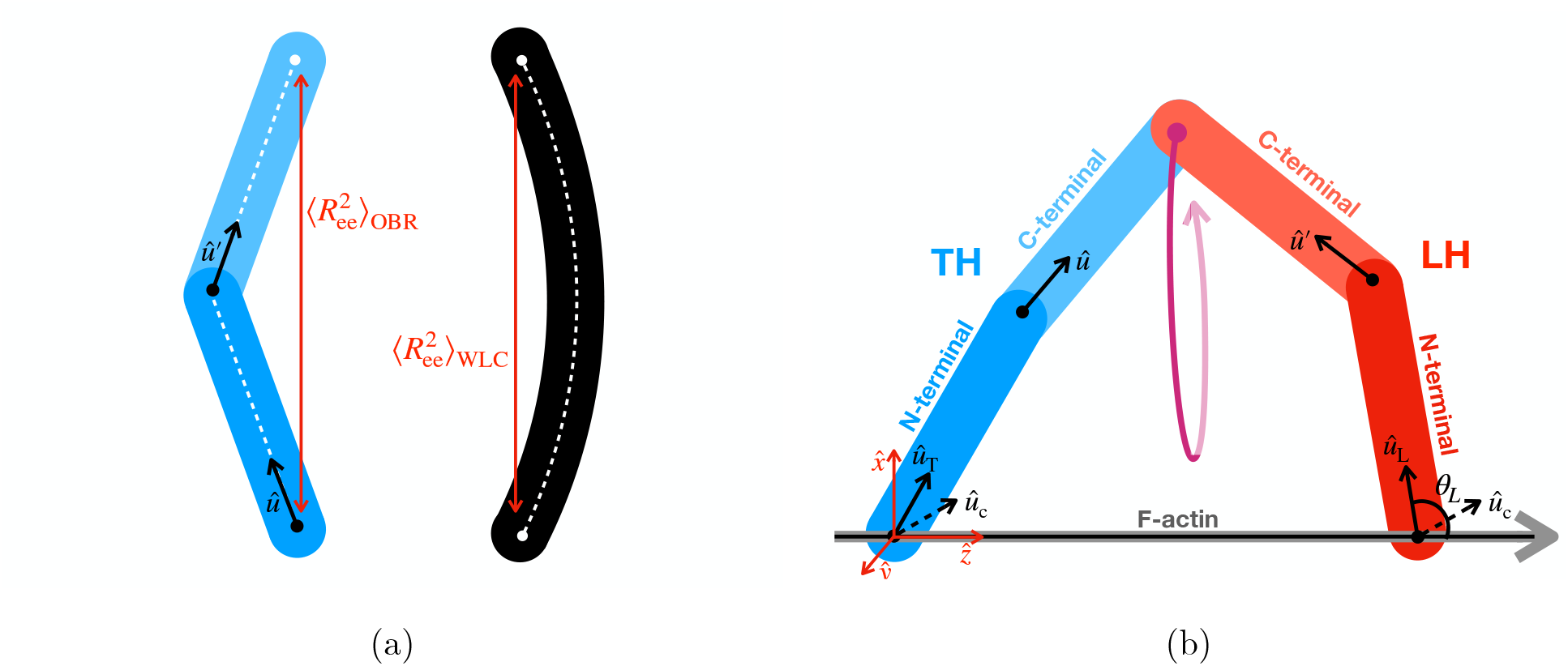
OBR model for the monomer and dimer. (a) Comparison of the OBR (left) and the WLC (right) models. The contour length (white, dashed lines) are the same. The hinge stiffness of the OBR is determined in order to recover the end-to-end distance (red lines) of a WLC with persistence length *l*_*p*_. (b) Dimeric construct. The TH lever arm is in blue and the LH lever arm is in red. The N-terminal rods are in darker color than the C-terminal segments. Given the orientation of the N-terminal segments, the C-terminal ones may rotate as shown by the purple line. The actin filament is shown in grey and black.

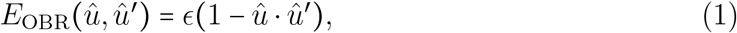

where *û* and *û*′ are the directions of the two rigid segments forming the OBR (see Fig. 1a). The length of the two rods is half of the lever arm contour length, and the stiffness of the hinge (*ϵ* in Eq. 1) is set to recover the average end-to-end distance of a worm-like chain (WLC) of persistence length *l*_*p*_ (see Fig. 1a and Appendix A). The OBR model, which allows us to perform analytic calculations, accounts for both the size and stiffness of the lever arm.

We term the two rigid rods constituting each lever arm as the N-terminal and C-terminal segments. The former is attached to the actin filament on one end (see Fig. 1b). The C-terminal segments of the LH and TH are joined together by a free joint (see Fig. 1b). The attachment of a motor to the actin filament induces a power stroke, which redirects the lever arm forward [see (11) for a detailed discussion]. The transition is assumed to have occurred already in the TH. It’s orientation with respect to the actin filament is restrained by a potential,

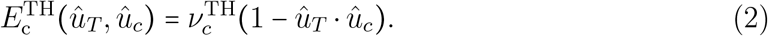

In the above equation 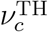 is the strength of the coupling between the TH lever arm and the motor domain, and *û*_*c*_ refers to the preferred orientation of the post-stroke conformation of the lever arm. The restraint on the LH is given by,

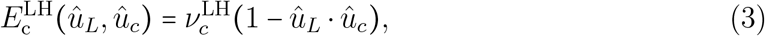

where the direction of the restraint (*û*_*c*_) is the same as for the TH. However, depending on the myosin model it may be possible that 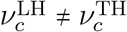 (see later). The statistical weight of a conformation, specified by the orientations *û*_*T*_ and *û*_*L*_ of the lever arms with respect to actin, is obtained by integrating over the conformations available for the C-terminal segments of the LH and TH OBRs – the joined ends may rotate as shown in Fig. 1b. As we show in Appendix A, the final result is,

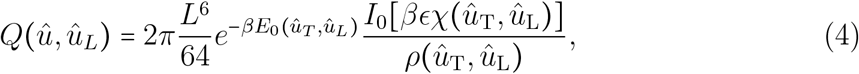

where *I*_0_ is a modified Bessel function of the first kind, and *E*_0_(*û*_*T*_, *û*_*L*_), *χ*(*û*_T_, *û*_L_), and *ρ* (*û*_T_, *û*_L_) are complicated functions of the orientations *û*_T_ and *û*_L_, and are given in Appendix A.

The probability density that the orientation of the LH is *û*_*L*_ is given by,

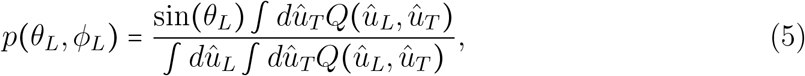

where *dû*_*L*_= *dθ*_*L*_*dϕ*_*L*_ sin (*θ*_*L*_) and *dû*_*T*_=*dθ*_*T*_ *dϕ*_*T*_ sin (*θ*_*T*_) are surface elements of a sphere of radius = 1. Note that the Jacobian for the LH angle is already incorporated in definition of *p*(*θ*_*L*_, *ϕ*_*L*_). The value of *p*(*θ*_*L*_, *ϕ*_*L*_)depends on: (i) *û*_*c*_, (ii) the stiffness of the lever arm, (iii) the contour length, and (iv) the strength of the orientation constraint in the LH. The average angle with the actin filament and the average orientation of the LH lever arm are given by,

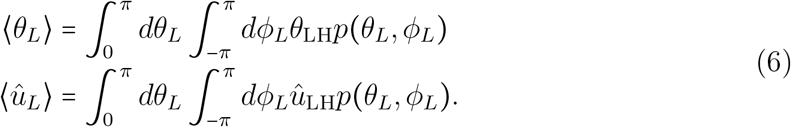

The integration over *û*_*T*_ and *û*_*L*_ was performed by Monte Carlo using the Metropolis algorithm (32), as detailed in Appendix B.

### Myosin V and Myosin VI

The parameters used for MV and MVI are in Table 1. The values were chosen on the basis of structural information and previous studies, as detailed in the caption in Table 1. Note that the choice of the strength of the TH constraint, 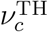, is identical in all models. This was set for MV in order to ensure that the LH lever arm orientation meets the value reported by Walker *et al*. (10). For the LH of MVI, we used *ν*_*c*_ = 0 to reflect the uncoupling of MVI LH from the motor domain (28).

The results in Fig. 2a-b highlight a dramatic difference: the orientation of the LH lever arm in MV is tightly constrained, whereas in MVI a broad range of angles are explored with nearly equal probability. Two possible features of MVI explain this drastically different behavior: (i) The lever arm of MVI arm is more flexible than MV (the value of *l*_*p*_ in MV is more than ten times greater than MVI, see Table 1); (ii) the lever arm of MVI uncouples from the motor domain. To isolate the contribution of these two features, we constructed two “chimeras”, one with the motor domain of MVI and MV lever arm, the other with the flexible lever arm of MVI that is coupled to the LH motor domain (as in the case of MV). Figures 2c-d show that changing either of these features results in dramatic changes of the explored conformations. For a quantitative analysis, we investigated the average angle of the LH lever arm measured from actin in the direction of forward walking (⟨*θ*_*L*_⟩) and the associated fluctuations of the LH lever arm, which are given by,

**Figure 2.**
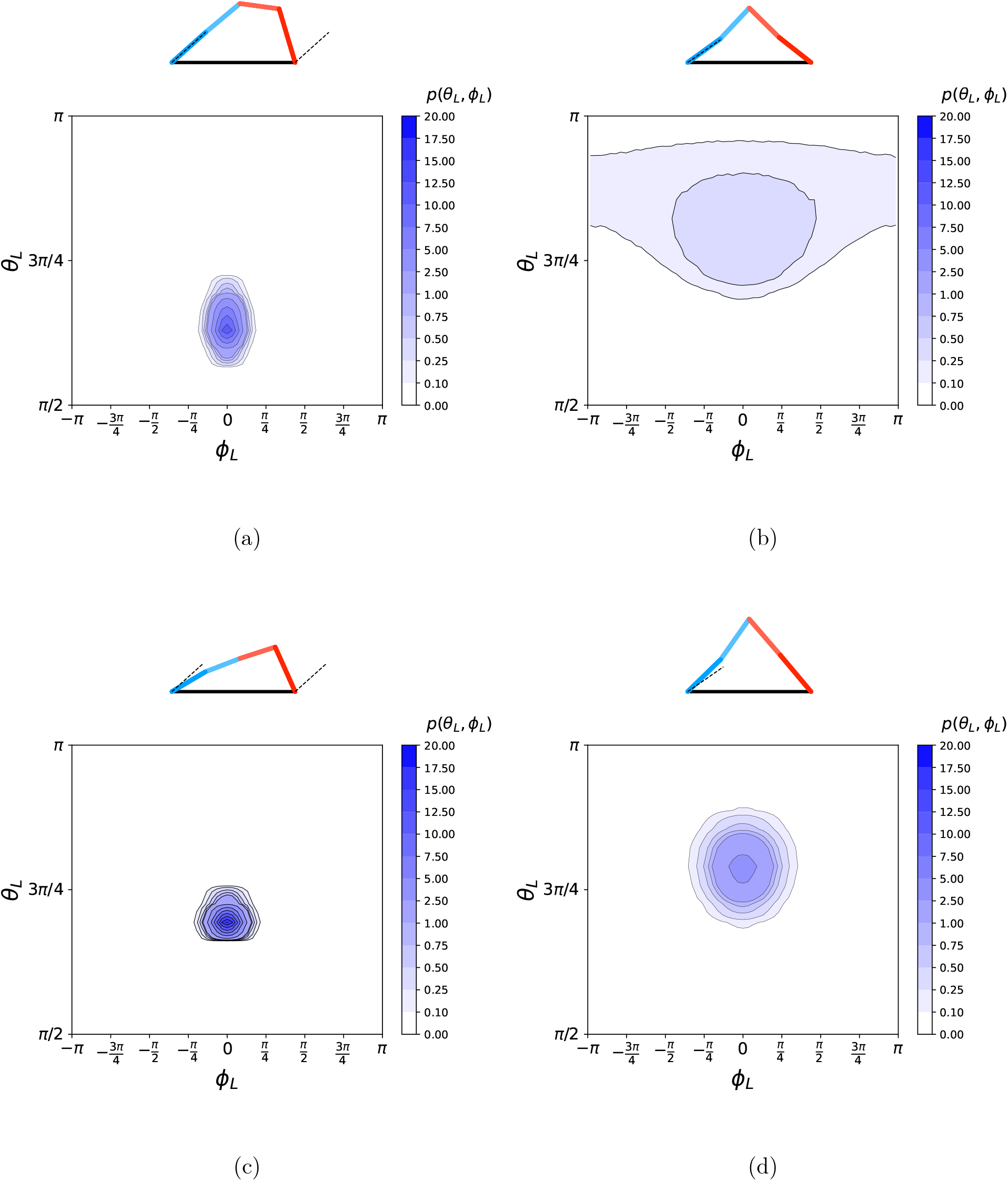
Theoretical predictions. In each figure, the sketch on top the panel illustrates the most likely conformation [the one that maximizes *Q* (*û*_*L*_, *û*_*T*_) (Eq. A19], and the dashed lines refer to the constrained imposed by the presence of actin, unless the LH is decoupled; the bottom panel is a contour plot showing *p* (*θ*_*L*_, *ϕ*_*L*_) (Eq. 5). (a) MV. The crossing points between the black, dashed lines refers to the average orientation of the LH reported by Walker et al. (10). (b) MVI. (c) Construct in which the motor domain is from MV and the lever arm from MVI. (d) Chimera with MVI motor domain and MV lever arm. Note that the color scale is the same in all figures.

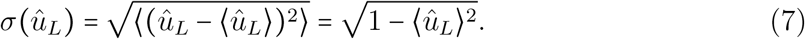

Table 2 shows that combining the MV motor domain with the MVI lever arm (MVmd/MVIla) affects predominantly the average angle with actin, making it somewhat larger compared to MV. The fluctuations of the LH are nearly unchanged. In contrast, replacing MVI lever arm with the stiffer MV counterpart (MVImd/MVla) slightly decreases the angle with actin while it nearly halves the size of fluctuations of the LH as compared to MVI.

**Table 2:** Results for the wild-type motors and the chimeras.

| parameter | MV | MVI | MVmd/MVlla | MVImd/MVla |
| --- | --- | --- | --- | --- |
| $\langle \theta_L \rangle$ | 2.01 | 2.61 | 2.19 | 2.47 |
| $\sigma(\hat{u}_L)$ | 0.17 | 0.50 | 0.16 | 0.27 |

By way of comparison to experiments, we note that Yildiz *et al*. (2) found that a Cy3 probe attached to the MVI lever arm fluctuates by *σ* ≈ 7 nm around its average position at physiological levels of ATP. In our model, given a probe bound to the N-terminal OBR of the LH at a distance *l* from the fixed point (attachment to the track), the variance of its location (*lσ* (*û*_*L*_)) is about 3 times larger in MVI than in MV, which is in qualitative agreement with experiments. Furthermore, in the MVI experiment (2) the probe was attached to the IQ motif-bound CaM, at a distance of *l*≈ 6.2 nm from the approximate location of the hinge of the rotation of MVI lever arm (around N785 (28)). In the OBR model, this gives *lσ* (*û*_*L*_) ≈ 3.1nm, that is smaller than the value reported in experiments (2). The discrepancy with experiments may be rationalized on the basis of a series of observations: (i) the experimental probe is off the long axis of MVI lever arm, (ii) further compliance in the motor domain might play a role by moving the approximate location of the hinge, (iii) the Cy3 location is estimated as the position of the alpha carbon of T146, not the source of signal, (iv) the actual MVI lever arm may be longer and more compliant than estimated here, and (v) sources of experimental inaccuracies may lead to an overestimate of the size of the fluctuations.

In addition, in a series of studies Goldman and coworkers investigated the different orientation of a lever-arm attached probe with respect to the actin filament in both the LH and TH of the dimer (23). The broader fluctuations observed in the leading head of MVI compared to the TH agree with the observations made by Yildiz (2) and hint at the existence of pliant region in MVI LH between the motor and the lever arm (25). In MV the distribution of orientations in the LH and TH are similar. As we show in the Appendix, we qualitatively recover these observations for both MV and MVI. A more quantitative comparison must account (i) for the precise orientation of the probe with respect to the lever arm, and (ii) for further elements of compliance that cannot be captured in the current model. However, we showed that coarse-grained models can recover these distributions successfully for MVI (28).

### Exploration of the Parameter Space

Experimentally it may be challenging to arbitrarily change the length and stiffness of the lever arm, or precisely control the degree of coupling between the lever arm and the motor domain. However, our theory could be used to continuously tune these parameters and explore the consequences of their modifications. Let us start from the parameters of MVI (see Table 1) and modify only the persistence length of the lever arm and the strength of the coupling with the motor domain. In other words, lever arm length and angle with actin are the same in the following paragraphs. Fig. 3 shows how the average angle between the LH and the actin filament (an observable that can be characterized using EM or AFM images of the processive dimer (4, 10, 23)) changes as a function of chain stiffness and strength of coupling to the LH. As the persistence length increases, the average angle slightly increases or remains nearly constant. In contrast, increasing the strength of coupling of the LH lever arm to the motor domain drastically reduces ⟨*θ*_LH_⟩. Increasing ⟨*θ*_LH_⟩ leads to conformations that resemble more the “telemark” stand (10); for stiff chains, this is a hallmark of the strong coupling between the LH motor domain and lever arm. Fig. 3a shows that, for 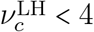, increasing the stiffness of the lever arm has little to no effect on the average angle of the LH. Note that for large stiffness at zero coupling to the lever arm we do not recover the limit of the chimera made of MVI motor domain and MV lever arm, because the length of the lever arm is fixed.

**Figure 3.**
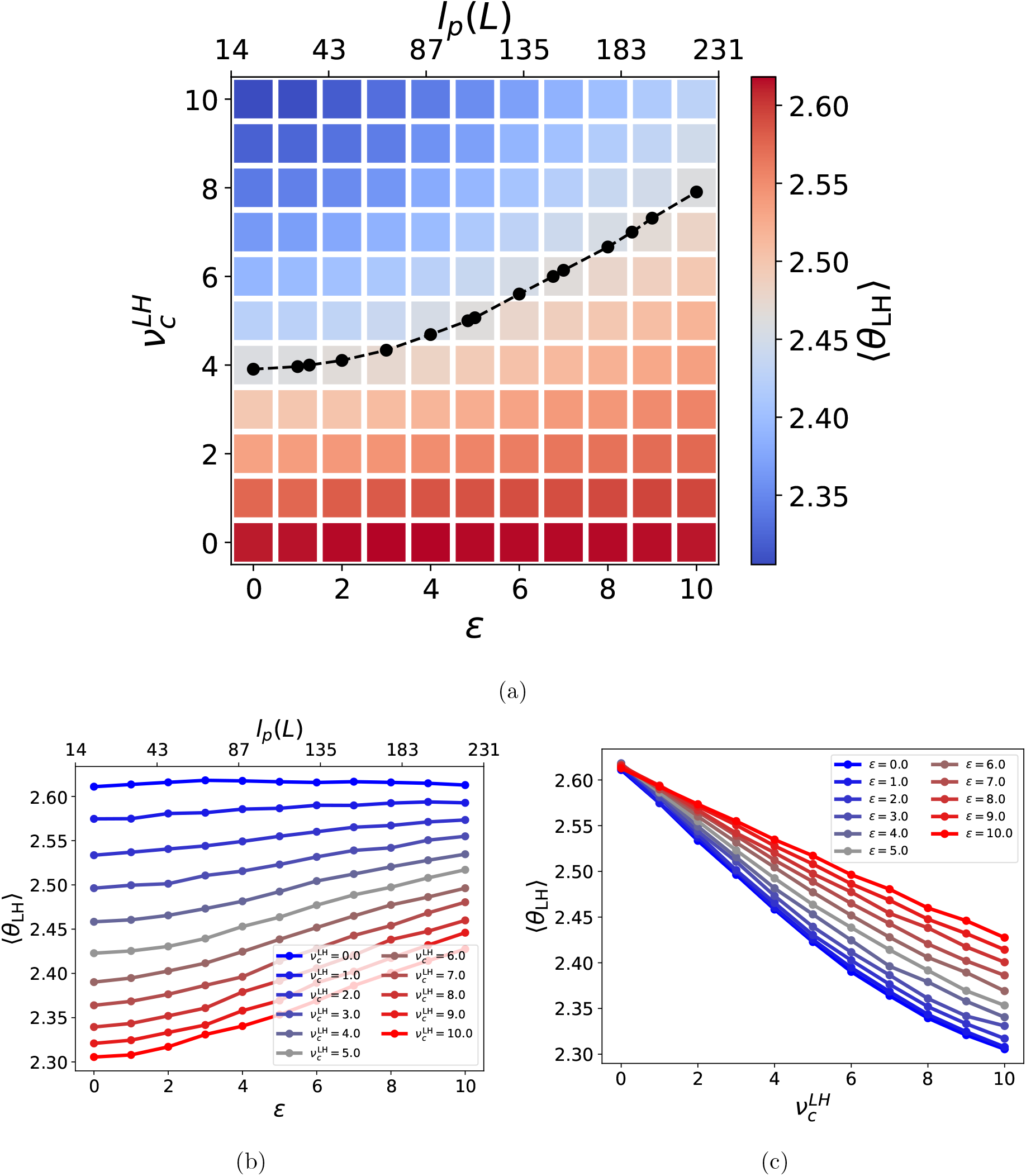
Dependence of the LH lever arm angle with the actin filament on the strength of coupling to the motor domain and the persistence length. (a) The colors indicate the value of ⟨*θ*_LH_⟩ – red is high, low is shown in blue. The x axis represent the stiffness (bottom) or persistence length (top) of the OBRs. The y axis is the strength of LH lever arm coupling to the motor domain. The black dots and line represent the locus of points for which ⟨*θ*_LH_⟩= [max (⟨*θ*_LH_⟩)+ min (⟨*θ*_LH_⟩]/2. The values were linearly extrapolated from computed data points. (b) ⟨*θ*_LH_⟩ as a function of the stiffness (bottom) or persistence length (top) of the lever arms. The colors refer to the strength of LH lever arm coupling. (c) ⟨*θ*_LH_⟩ as a function of *ν*_*c*_. The color refers to the stiffness of the chain.

Figure 4 shows the effect of chain stiffness and strength of lever arm coupling to the LH on the fluctuations of the LH lever arm (*σ* (*û*_*L*_)), which can be characterized experimentally by using fluorescent probes attached to the lever arm (2, 23). The fluctuations are larger when *ϵ* and 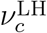 are both small, and are progressively reduced as either of the two is increased. Interestingly, this does not occur monotonically: an increase in 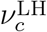 may lead to a small increase of *σ* (*û*_*L*_), as shown in Fig. 4c. This is explained by performing a Taylor expansion of *σ* (*û*_*L*_) for small fluctuations (Δ*θ* and Δ*ϕ*) in the vicinity of the average angle orientation (indicated with an overline, e.g. 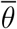). The result is that 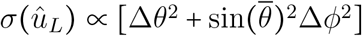, which implies that an increase of the coupling with the LH lever arm would increase 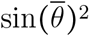 without affecting the size of the fluctuations, resulting in the amplification *σ* (*û*_*L*_). Finally, it is notable (Fig. 4a) that if the lever arm is sufficiently stiff (*l*_*p*_ ⪆ 135 nm) increasing the strength of the coupling to the motor domain has essentially no effect on the size of the fluctuations: they remain small. Similarly, if the coupling is sufficiently strong 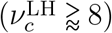, making the lever arm stiffer does not significantly increase the fluctuation of the LH.

**Figure 4.**
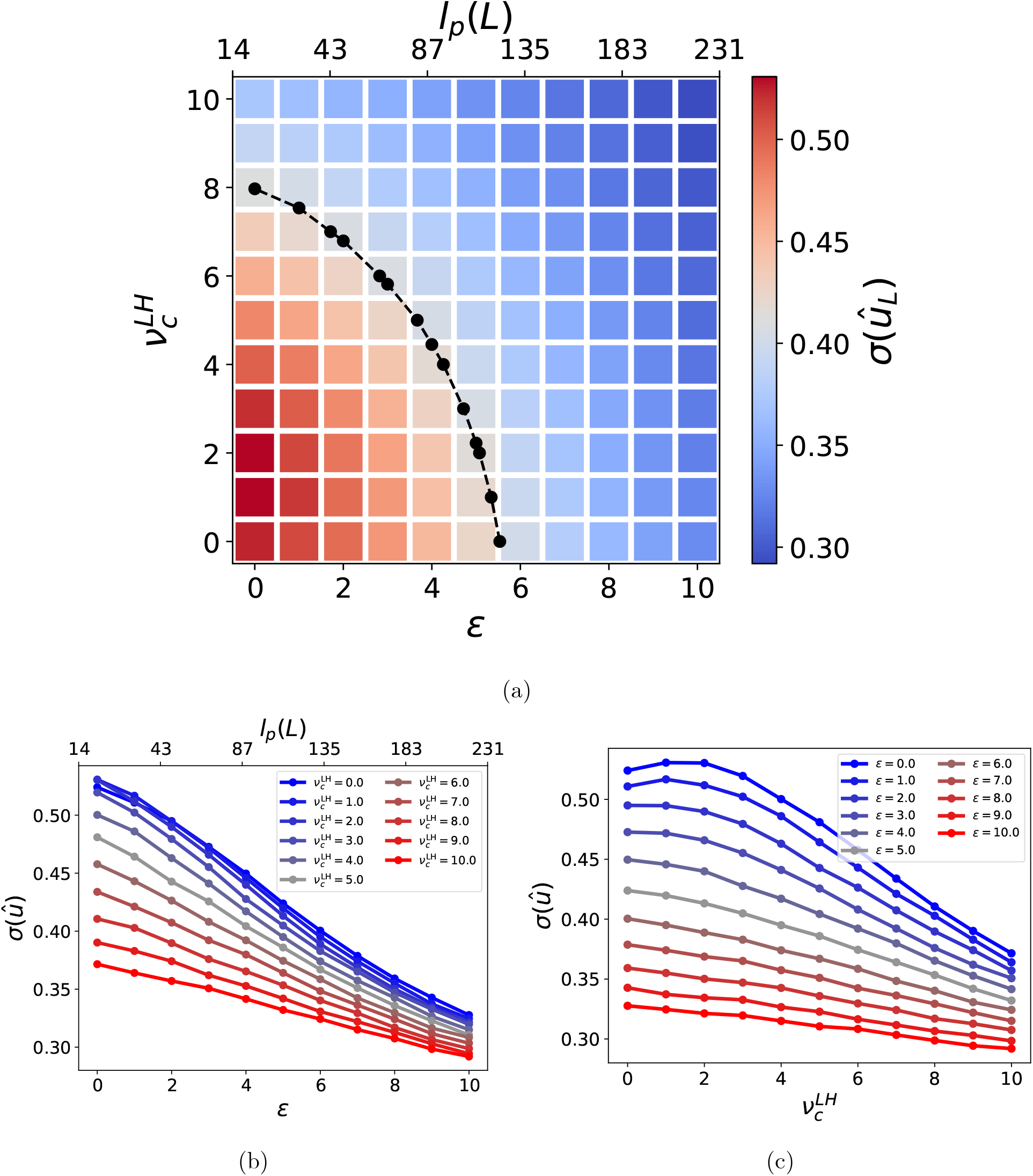
Dependence of the conformational flexibility of the LH on the strength of coupling to the motor domain and the persistence length. Panels and color codes are the same as in Fig. 3, but refer to *σ*(*û*_*L*_).

## Discussion

We developed a simple model in which the geometry of the two-head bound conformation of myosin dimer is described by (i) the average step-size of the motor (*d*), (ii) the contour length of the lever arm (*L*), (iii) the stiffness of the lever arm (*ϵ*), (iv) the favored orientation of the lever arm with respect to the actin filament in the actin-bound state (*û*_*c*_), and (v) the stiffness of the restraint of the lever arm orientation around its preferred orientation (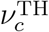 and 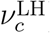). Although we believe this to be a minimal set of parameters, there are some important features that are neglected. For instance, (i) the step size distribution of the motor is not fixed, therefore *d* changes in different conformations, (ii) this also implies that the azimuthal angle between LH and TH lever arm can be non-zero; (iii) the joint between the LH and TH lever arms might not be free (see Andrecka *et al* (33) and Hathcock *et al* (30)); (iv) the interaction between the motor domain and the LH lever arm might restrain some conformations from occurring even in the uncoupled state.

We studied the conformational flexibility and the average angle of the LH lever arm with respect to the motor domain in four constructs: wild-type MV and MVI, and chimeras with the motor domain of MV (MV) combined with MVI (MV) lever arm. It is possible to create these constructs (see for instance Park *et al*. (14)), and therefore the changes that we observed (see Table 2) constitute predictions that amenable to experimental test. The static theory describing the fluctuations of the lever arm geometries cannot be used to account for the stepping kinetics. Nevertheless, the histograms accounting for the fluctuations in *θ* (Fig. A3) associated with the LH and TH provides insights into the differences in the lever arm rotation between the motors. The differences in the location of the peaks in Fig. A3 is an estimate of the the angle by which the lever arm in the two motors rotate. In MV (MVI) the lever rotates by ≈90° (≈135°). Several studies (13, 25, 27, 34) indicated that MVI undergoes a large rotation of the lever arm, possibly up to ≈180°. Our results do not get to such large values, but do indicate that the MVI stroke is larger than MV (see Fig. A3).

To garner further understanding of the interplay between stiffness of the chain and strength of coupling between LH lever arm and motor domain, we explore systematically how these two features affect the average angle with respect to the actin filament, ⟨*θ*_*L*_⟩, and a measure of the variance of its orientation, *σ* (*û*_*L*_). Both of these quantities are experimentally measurable by attaching fluorescent probes to the lever arm. We set *L, d, û*_*c*_, and 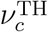 to be equal to their values for MVI and make the following observations.

a. If the coupling between LH lever arm and motor domain is weak 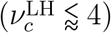 the angle with the actin filament measured from the preferred direction of movement is large, irrespective of the stiffness of the lever arm (see Fig. 3a).
b. If the lever arm is stiff (*l*_*p*_ ⪆ 130 nm), the fluctuations of the LH lever arm are suppressed, regardless of the strength of the coupling between LH motor domain and lever arm (see Fig. 4a). Analogously, if the coupling between motor domain and LH lever arm is large enough 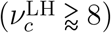 the fluctuations are suppressed irrespective of lever arm stiffness (see Fig. 4a).
c. The average angle ⟨*θ*_*L*_⟩ results from a balance of stiffness of the lever arm, which tends to decrease it, and strength of the coupling with the motor domain, whose effect is to reduce the angle and promote a “telemark”-like conformation. The antagonistic effect is highlighted by the positive slope of the black curve in Fig. 3a.
d. In contrast, increasing either 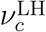 or *ϵ* suppresses the fluctuations of the LH lever arm, as indicated by the negative slope of the black curve in Fig. 4a.

Our results suggest then some rules that are useful in interpreting AFM, EM, and Total Internal Reflection Fluorescence (TIRF) experiments reporting on the structure and conformational flexibility of the LH lever arm of a dimer. By comparing to our model, it may be possible to ascertain whether the information is sufficient to draw conclusion on the lever arm stiffness and the strength of coupling between lever arm and motor domain. This could help in understanding processive myosin motors, as well as the design on new molecular machines.

## Appendix A Model

### Once-broken Rod Model

We describe the lever arm of myosin motors as two rigid rods connected by a restrained hinge (Once-broken Rod, OBR, see Fig. 1a). The energy function of the OBR is given by Eq. 1. The mean-squared end-to-end distance of the OBR is given by,

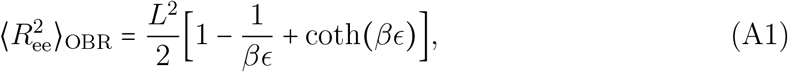

where *L* is the contour length of the polymer (the sum of the lengths of the two rigid rods) and *β* ^−1^= *k*_*B*_*T*. The model has two parameters: the contour length (*L*) and the chain stiffness (*ϵ*). In the limit of large *βϵ*, Eq. A1 reduces to 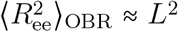. A better description of the lever arm is the worm-like chain (WLC), in which the flexibility of the polymer is distributed across the entire length of the polymer. The mean-squared end-to-end distance of the WLC is (37),

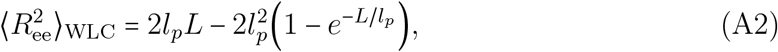

where *l*_*p*_ is the persistence length. Experiments can estimate both the contour length and the persistence length of the lever arm. The OBR and the WLC models can be mapped onto each other in some limits. We search for the value of *ϵ* to ensure that 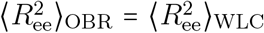. Note that 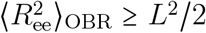, where the equality holds for *ϵ* = 0. If the end-to-end distance of the WLC is smaller than *L*^2^ 2 this procedure fails. However, in our case the smallest persistence length that can be mapped onto the OBR model is smaller than the persistence length of CaM-bound lever arm and of the microtubule.

#### Model for actin-bound LH and TH

Let two identical lever arms, modeled as OBRs, represent the LH and TH of a myosin dimer bound to the actin filament. F-actin is parallel to the 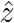 axis; therefore one end of the TH OBR is located at the origin of the axis, and one end of the LH OBR is at 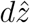, where *d* ≈ 36 nm corresponding to the step-size of myosin and half-pitch of F-actin. The other ends of both OBRs are connected with a free joint (see Fig. A1).

The TH and LH are subject to the following potential (see Fig. A1), which restrains the first segment of each lever arm,

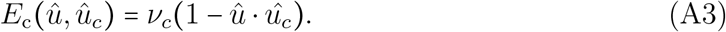

The constraint angle is taken to be the equilibrium orientation for the post-stroke conformation. Therefore, the overall energy function is,

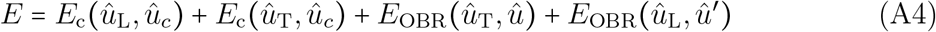

The first two segments of the lever arm are connected by a vector 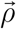 given by,

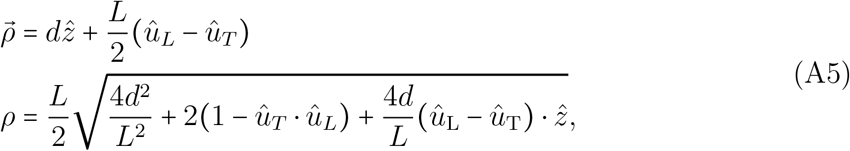

where *d* is the projection of 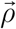 along the 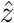axis and *ρ* is the length of 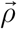. The segments of the lever arms which are restrained by the interaction with the filament are joined together and can rotate around the vector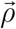.

There are three possible scenarios: (i) 0 *ρ* = *L*, which is depicted in Fig. A1; (ii) ∣ *ρ*∣ = *L*, in which case no rotation around 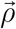 is possible; (iii) *ρ* = 0, occurring when the two rigid initial segments of the lever arms end at the same location.

#### Partition Function when 0< ∣*ρ*∣< *L*

The model can be described using 5 beads represented as black dots in Fig. A1. Given 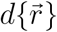 as the phase space volume of the five beads, the partition function is given by,

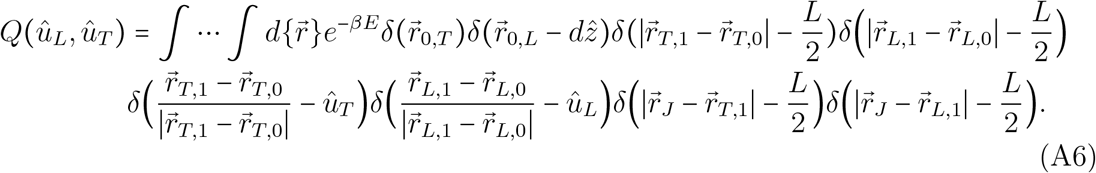

The first two delta functions fix the location of the lever arms along F-actin, the next four dictate the orientation and size of the first rods of both OBRs, the remaining two state that the size of the second rods of the OBRs is *L*/ 2. After carrying out the integrals we are left with,

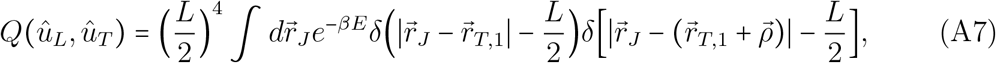

where the dependence on *û* _*L*_ and *û* _*T*_ is inside 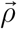 (see Eq. A5). To perform this integral we adopt the reference system, 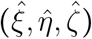 (see Fig. A1), with 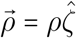, and 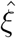 oriented in the direction of the component of *û*_T_ + *û*_L_ which is perpendicular to 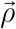 (see Fig. A1), so that,

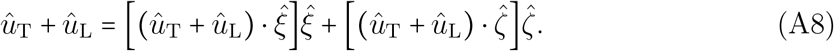

Using this new reference system, the partition function becomes,

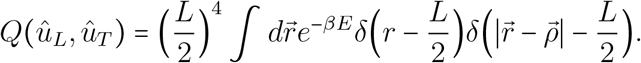

The second delta function can be reformulated as an angular constraint with some ma-nipulations,

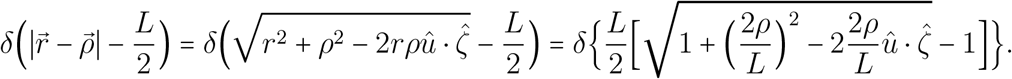

For the last equality we used 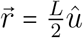. Let us define,

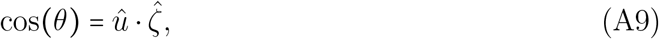

and using the properties of the delta distribution we obtain,

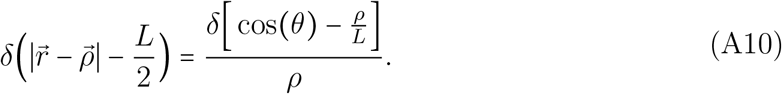

The partition function becomes,

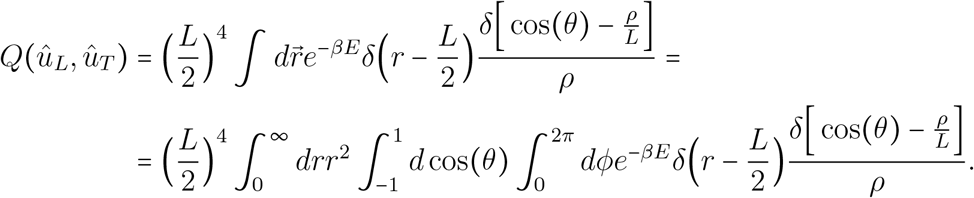

Upon carrying out the integration over the delta functions, we obtain,

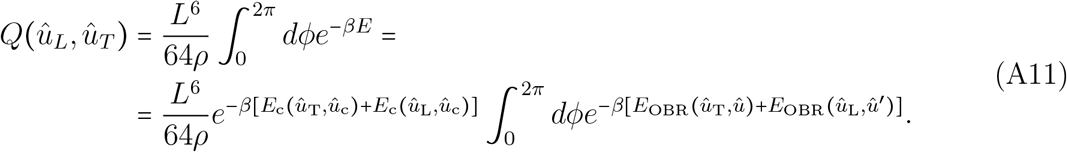

Note that, because of the constraints we can write,

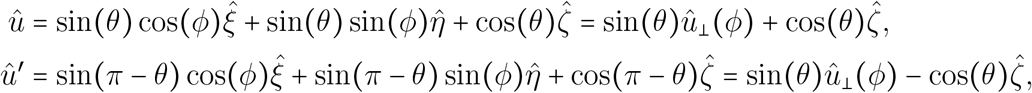

where 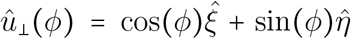 is a unit vector in the 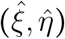 plane and carries the dependence on *ϕ*. Replacing cos(*θ*) = *ρ*/*L*, and recalling that 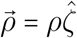, we can now write,

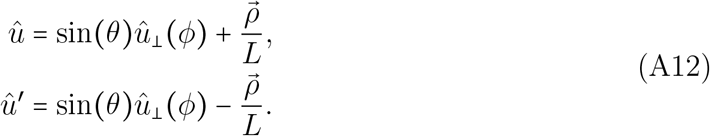

Using these expressions we may simplify the contribution to the total energy function in Eq. A4 of the OBR terms,

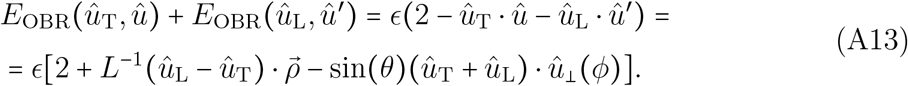

Using Eq. A5 we can rewrite,

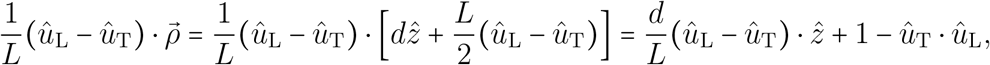

so that,

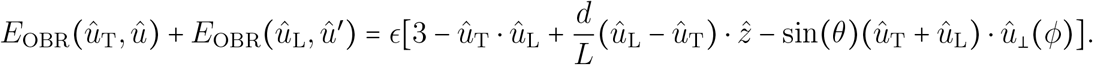

Using Eq. A8 we obtain,

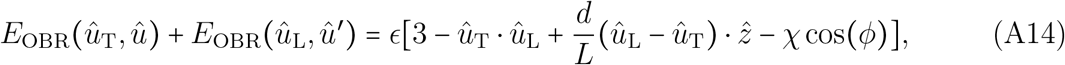

where,

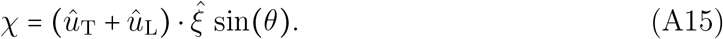

Finally, we compute 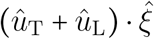 as,

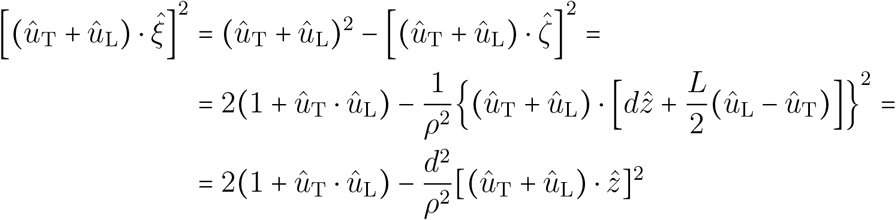

so that finally,

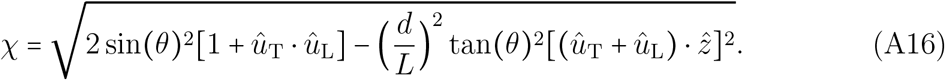

We now rewrite the energy function as,

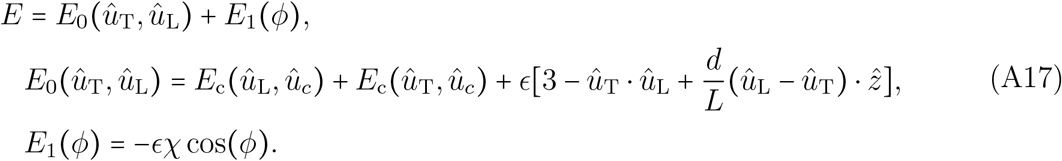

The partition function in Eq. A11 can then be rewritten as,

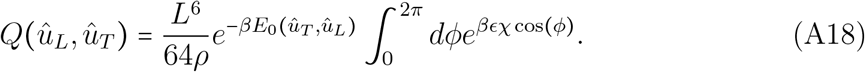

Given that 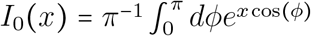 is the modified Bessel function of the first kind, we can write.

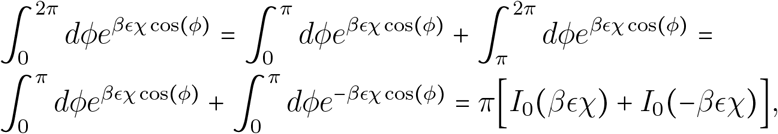

It can be shown that *I*_0_(*x*) is an even function, *I*_0_(*x*) = *I*_0_(− *x)*, which can be shown by replacing the integration variable *ϕ* = *π* − *ψ*:

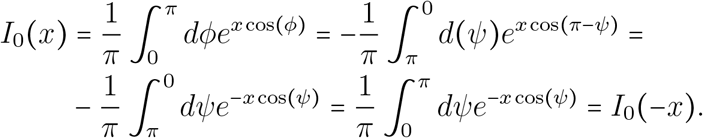

Thus, we finally obtain,

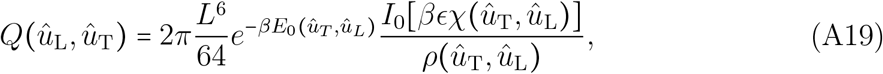

#### Partition Function when ∣*ρ*∣≥ *L*

If ∣*ρ*∣ > *L*, then it is not possible to connect the lever arms, and the corresponding conformation has probability 0. We assign the same probability to the case *ρ*=*L*, for which there is a single allowed conformation with the second rod of both lever arms are connected to each other and with an angle between them of *π*. Other choices could be physically sensible; Eq. A19 has a limit for *ρ*=*L*, and we could take the value in Eq. A19 and exclude the contribution from the integral over the azymuthal angle (2*πI*_0_(*βϵχ*)).

#### Partition Function when ∣*ρ*∣ = 0

Clearly we run into some difficulties when taking the limit for *ρ* → 0 in Eq. A19. However, in order for ∣*ρ*∣ = 0 we need *L* cos(*θ* _*L*_) = *d*, but in the cases that we consider *d* > *L*, therefore ∣*ρ*∣ ≠ 0.

#### Parameters of the MV and MVI Lever Arm Models

The MV lever arm was estimated from the structure resolved by Liu *et al*. (35) (see Fig. A2a). To estimate the length of the lever arm we assembled three domains from three different structures. The alignment is in Fig. A2b.

## Appendix B Numerical Integration

Computing the probability 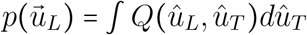 requires an integral over the surface of a sphere of *Q* (*û*_*L*_, *û*_*T*_). We decided to perform the integral using a Monte Carlo approach highlighted as follows:

1. Start with a conformation of the dimer with weight *w*_*i*_*=*0, given by {*û*_*T*_, *û*_*L*_};
2. Randomly select the angle to update; the new conformation is 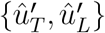
3. Reject the conformation if ∣*ρ*∣ ≥ *L* or *ρ* = 0: we update *w*_*i*_ = *w*_*i*_ + 1 and go back to step 2;
4. If 0< ∣*ρ*∣< *L*, we compute 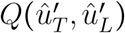;
5. Accept the conformation with probability 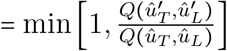 (32);
6. If the conformation is accepted, *i* = *i* + 1, *w*_*i*_ = 1, and we go back to step 2;
7. If the conformation is rejected, *w*_*i*_= *w*_*i*_ +1, and we go back to step 2.

The process is repeated 10^7^ times; the points are sampled from ∝*Q* (*û*_*T*_, *û*_*L*_), and the average of any observable *O*(*û*_*T*_, *û*_*L*_) is,

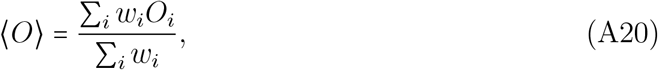

where the sum is over all the conformations sampled.

It is important to highlight that when we draw the new set of angles, the following procedure is adopted. Let *u* be a random number sampled from a uniform distribution between 0 and 1; if we update a *ϕ* angle, then the new angle is *ϕ =* 2*πu* −*π*. Instead of updating a *θ* angle, we update the cosine of that angle, that is Θ cos = (*θ*) 2*u* −1; the angle will then be *θ =* arccos (Θ). This procedure includes in the sampling the Jacobian for spherical coordinates (*dθ* sin(*θ*) = *d* cos(*θ*)), and we do not need to include it in *Q*(*û*_*T*_, *û*_*L*_).

## Acknowledgments

This work was supported by a grant from the National Science Foundation (CHE 2320256) and the Welch Foundation through the Collie-Welch Chair (F-0019).

---

**Figure A1:**
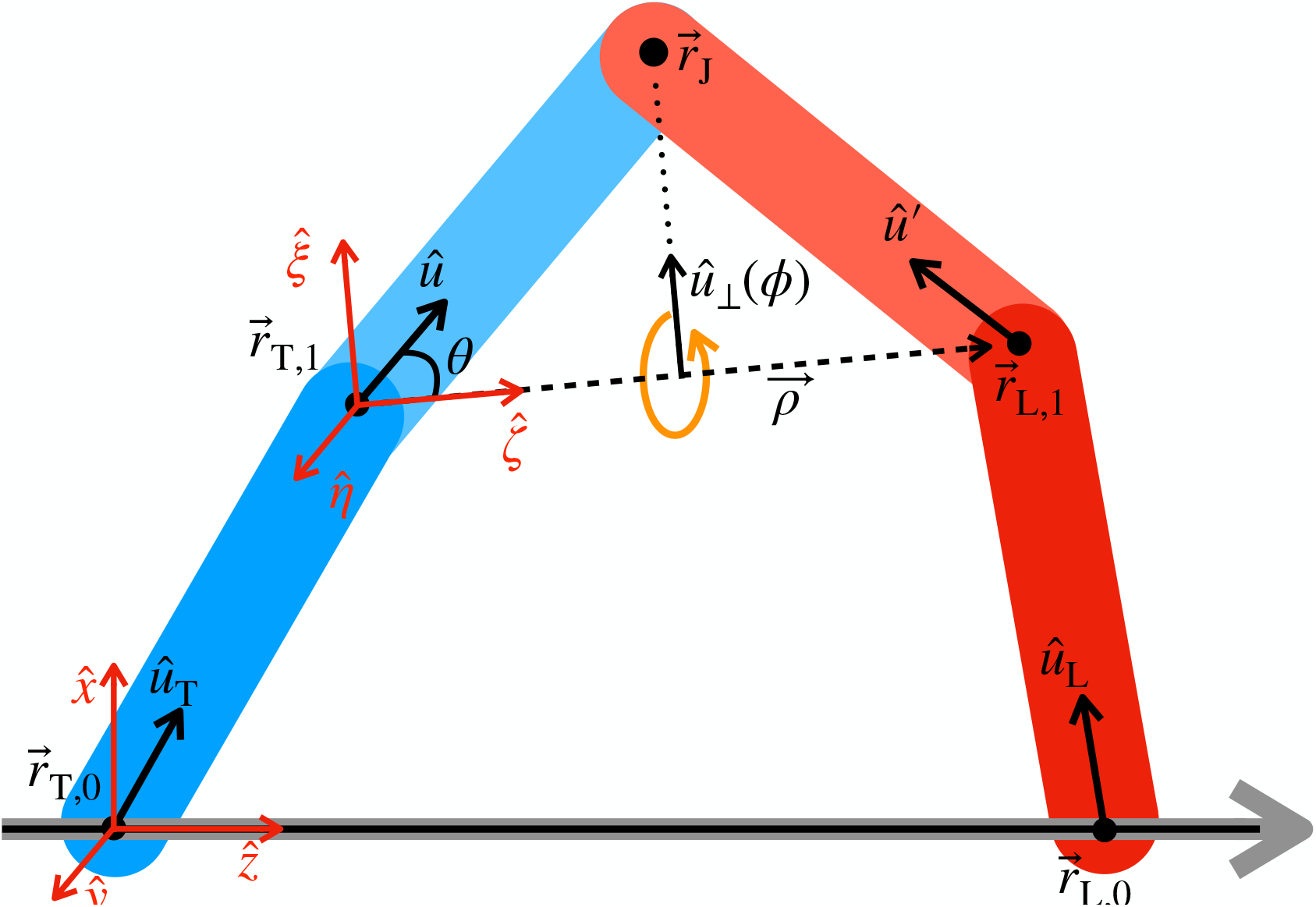
OBR model for the dimer. The TH (LH) lever arm is in blue (red). The actin filament is shown in gray and black. The vector *ρ* is in dashed, black lines, and the dotted vector has the direction of *û*_⊥_(*ϕ*), corresponding to the component of *û* (and *û*^′^) which is perpendicular to 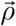. Two reference systems are shown in red: one, 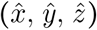 has 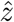 parallel to the actin filament, and the vector *û*_T_ is in the 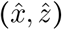 plane. The second reference system, 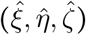has 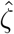 parallel to 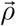, and 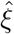 parallel to the component of *û*_T_ *û*_L_ that is perpendicular to 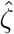. The statistics of the OBR can be described using the 5 beads depicted as black dots.

**Figure A2:**
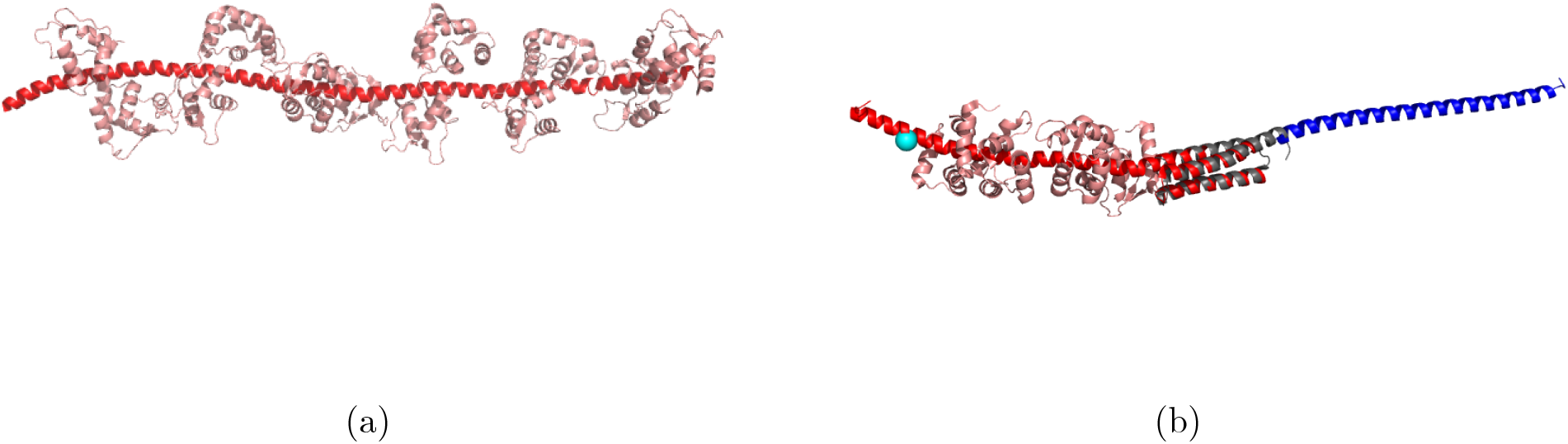
Model for MV and MVI lever arms. (a) MV lever arm, from PDBID: 2DFS (35). Residues 755 and 909 in red, and the six bound calmodulins (CaMs) are in pink. (b) For MVI, we combined three elements: the ins2 and CaM binding domain from PDBID: 3GN4 (red, white pink CaMs) (16); gray shows the proximal tail (PT) from PDBID: 2LD3; the blue Single-alpha-helix (SAH) medial tail (MT) is from PDBID: 6OBI (22). The alignment of the PT to the MT is somewhat arbitrary: the residue overlap between the two groups is too small. The cyan sphere indicates N785, a putative hinge for the lever arm (28). Overall, the length of the lever arm from N785 to the end of the MT is approximately 21 nm. The alignment was performed using PyMol.

**Figure A3:**
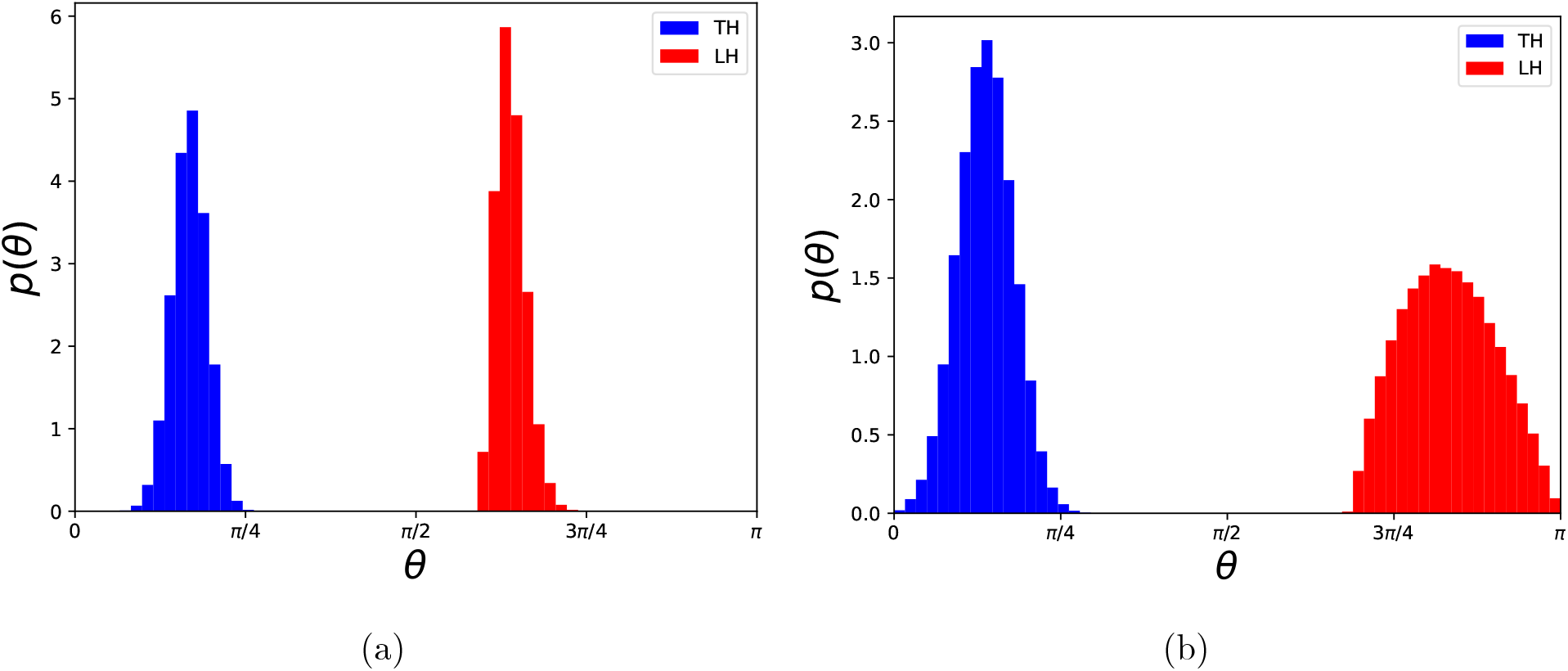
Distributions of the LH and TH angle with respect to the actin filament. (a) MV, the TH and LH are shown in blue and red, respectively. (b) MVI, with the same color code as (a).

## References

[1] Ahmet Yildiz, Joseph N. Forkey, Sean A. McKinney, Taekjip Ha, Yale E. Goldman, and Paul R. Selvin. Myosin V Walks Hand-Over-Hand: Single Fluorophore Imaging with 1.5-nm Localization. Science, 300(5628):2061–2065, 2003.

[2] Ahmet Yildiz, Hyokeun Park, Dan Safer, Zhaohui Yang, Li-Qiong Chen, Paul R. Selvin, and H. Lee Sweeney. Myosin VI Steps via a Hand-over-Hand Mechanism with Its Lever Arm Undergoing Fluctuations when Attached to Actin. Journal of Biological Chemistry, 279(36):37223–37226, 2004.

[3] Zeynep Ökten, L Stirling Churchman, Ronald S Rock, and James A Spudich. Myosin VI walks hand-over-hand along actin. Nature Structural & Molecular Biology, 11(9):884–887, 2004.

[4] Noriyuki Kodera, Daisuke Yamamoto, Ryoki Ishikawa, and Toshio Ando. Video imaging of walking myosin V by high-speed atomic force microscopy. Nature, 468 (7320):72–76, 2010.

[5] So Nishikawa, Ikuo Arimoto, Keigo Ikezaki, Mitsuhiro Sugawa, Hiroshi Ueno, Tomotaka Komori, Atsuko H. Iwane, and Toshio Yanagida. Switch between Large Hand-Over-Hand and Small Inchworm-like Steps in Myosin VI. Cell, 142(6):879– 888, 2010.

[6] Mauro L Mugnai, Changbong Hyeon, Michael Hinczewski, and D Thirumalai. Theoretical perspectives on biological machines. Reviews of Modern Physics, 92(2):025001, 2020.

[7] Anatoly B Kolomeisky and Michael E Fisher. Molecular motors: a theorist’s perspective. Annu. Rev. Phys. Chem., 58:675–695, 2007.

[8] Enrique M. De La Cruz, Amber L. Wells, Steven S. Rosenfeld, E. Michael Ostap, and H. Lee Sweeney. The kinetic mechanism of myosin V. Proceedings of the National Academy of Sciences, 96(24):13726–13731, 1999.

[9] Enrique M. De La Cruz, E. Michael Ostap, and H. Lee Sweeney. Kinetic Mechanism and Regulation of Myosin VI. Journal of Biological Chemistry, 276(34):32373–32381, 2001.

[10] Matthew L. Walker, Stan A. Burgess, James R. Sellers, Fei Wang, John A. Hammer, John Trinick, and Peter J. Knight. Two-headed binding of a processive myosin to F-actin. Nature, 405(6788):804–807, 2000.

[11] Michael Hinczewski, Riina Tehver, and D Thirumalai. Design principles governing the motility of myosin V. Proceedings of the National Academy of Sciences, 110(43):E4059–E4068, 2013.

[12] Amber L. Wells, Abel W. Lin, Li-Qiong Chen, Daniel Safer, Shane M. Cain, Tama Hasson, Bridget O. Carragher, Ronald A. Milligan, and H. Lee Sweeney. Myosin VI is an actin-based motor that moves backwards. Nature, 401(6752):505–508, 1999.

[13] Zev Bryant, David Altman, and James A. Spudich. The power stroke of myosin VI and the basis of reverse directionality. Proceedings of the National Academy of Sciences, 104(3):772–777, 2007.

[14] Hyokeun Park, Anna Li, Li-Qiong Chen, Anne Houdusse, Paul R. Selvin, and H. Lee Sweeney. The unique insert at the end of the myosin VI motor is the sole determinant of directionality. Proceedings of the National Academy of Sciences, 104(3):778–783, 2007.

[15] Benjamin J Spink, Sivaraj Sivaramakrishnan, Jan Lipfert, Sebastian Doniach, and James A Spudich. Long single α-helical tail domains bridge the gap between structure and function of myosin VI. Nature Structural & Molecular Biology, 15(6):591–597, 2008.

[16] Monalisa Mukherjea, Paola Llinas, HyeongJun Kim, Mirko Travaglia, Daniel Safer, Julie Ménétrey, Clara Franzini-Armstrong, Paul R. Selvin, Anne Houdusse, and H. Lee Sweeney. Myosin VI Dimerization Triggers an Unfolding of a Three-Helix Bundle in Order to Extend Its Reach. Molecular Cell, 35(3):305–315, 2009.

[17] D. Thirumalai and Zhechun Zhang. Myosin VI: How Do Charged Tails Exert Control? Structure, 18(11):1393–1394, 2010.

[18] HyeongJun Kim, Jen Hsin, Yanxin Liu, Paul R. Selvin, and Klaus Schulten. Formation of Salt Bridges Mediates Internal Dimerization of Myosin VI Medial Tail Domain. Structure, 18(11):1443–1449, 2010.

[19] Yanxin Liu, Jen Hsin, HyeongJun Kim, Paul R. Selvin, and Klaus Schulten. Extension of a Three-Helix Bundle Domain of Myosin VI and Key Role of Calmodulins. Biophysical Journal, 100(12):2964–2973, 2011.

[20] Monalisa Mukherjea, M. Yusuf Ali, Carlos Kikuti, Daniel Safer, Zhaohui Yang, Helena Sirkia, Virginie Ropars, Anne Houdusse, David M. Warshaw, and H. Lee Sweeney. Myosin VI Must Dimerize and Deploy Its Unusual Lever Arm in Order to Perform Its Cellular Roles. Cell Reports, 8(5):1522–1532, 2014.

[21] Sivaraj Sivaramakrishnan, Benjamin J. Spink, Adelene Y. L. Sim, Sebastian Doniach, and James A. Spudich. Dynamic charge interactions create surprising rigidity in the ER/K α-helical protein motif. Proceedings of the National Academy of Sciences, 105 (36):13356–13361, 2008.

[22] C. Ashley Barnes, Yang Shen, Jinfa Ying, Yasuharu Takagi, Dennis A. Torchia, James R. Sellers, and Ad Bax. Remarkable Rigidity of the Single α-Helical Domain of Myosin-VI As Revealed by NMR Spectroscopy. Journal of the American Chemical Society, 141(22):9004–9017, 2019.

[23] Yujie Sun and Yale E. Goldman. Lever-Arm Mechanics of Processive Myosins. Biophysical Journal, 101(1):1–11, 2011.

[24] Giovanni Cappello, Paolo Pierobon, Clémentine Symonds, Lorenzo Busoni, J. Christof, M. Gebhardt, Matthias Rief, and Jacques Prost. Myosin v stepping mechanism. Proceedings of the National Academy of Sciences, 104(39):15328–15333, 2007.

[25] Yujie Sun, Harry W Schroeder, John F Beausang, Kazuaki Homma, Mitsuo Ikebe, and Yale E Goldman. Myosin vi walks “wiggly” on actin with large and variable tilting. Molecular cell, 28(6):954–964, 2007.

[26] Jeff G. Reifenberger, Erdal Toprak, HyeongJun Kim, Dan Safer, H. Lee Sweeney, and Paul R. Selvin. Myosin VI undergoes a 180° power stroke implying an uncoupling of the front lever arm. Proceedings of the National Academy of Sciences, 106(43):18255–18260, 2009.

[27] Julie Ménétrey, Tatiana Isabet, Virginie Ropars, Monalisa Mukherjea, Olena Pylypenko, Xiaoyan Liu, Javier Perez, Patrice Vachette, H. Lee Sweeney, and Anne M. Houdusse. Processive Steps in the Reverse Direction Require Uncoupling of the Lead Head Lever Arm of Myosin VI. Molecular Cell, 48(1):75–86, 2012.

[28] Mauro L Mugnai and D Thirumalai. Kinematics of the lever arm swing in myosin VI. PNAS, 114(22):E4389–E4398, 2017.

[29] Erin M. Craig and Heiner Linke. Mechanochemical model for myosin V. Proceedings of the National Academy of Sciences, 106(43):18261–18266, 2009.

[30] David Hathcock, Riina Tehver, Michael Hinczewski, and D Thirumalai. Myosin V executes steps of variable length via structurally constrained diffusion. Elife, 9:e51569, 2020.

[31] Ganhui Lan and Sean X. Sun. Flexible Light-Chain and Helical Structure of FActin Explain the Movement and Step Size of Myosin-VI. Biophysical Journal, 91 (11):4002–4013, 2006.

[32] Nicholas Metropolis, Arianna W. Rosenbluth, Marshall N. Rosenbluth, Augusta H. Teller, and Edward Teller. Equation of state calculations by fast computing machines. The Journal of Chemical Physics, 21(6):1087–1092, 1953.

[33] Joanna Andrecka, Jaime Ortega Arroyo, Yasuharu Takagi, Gabrielle de Wit, Adam Fineberg, Lachlan MacKinnon, Gavin Young, James R Sellers, and Philipp Kukura. Structural dynamics of myosin 5 during processive motion revealed by interferometric scattering microscopy. eLife, 4:e05413, 2015.

[34] Ida Lister, Stephan Schmitz, Matthew Walker, John Trinick, Folma Buss, Claudia Veigel, and John Kendrick-Jones. A monomeric myosin vi with a large working stroke. The EMBO Journal, 23(8):1729–1738, 2004.

[35] Jun Liu, Dianne W. Taylor, Elena B. Krementsova, Kathleen M. Trybus, and Kenneth A. Taylor. Three-dimensional structure of the myosin V inhibited state by cryoelectron tomography. Nature, 442(7099):208–211, 2006.

[36] Pinar S Gurel, Laura Y Kim, Paul V Ruijgrok, Tosan Omabegho, Zev Bryant, and Gregory M Alushin. Cryo-EM structures reveal specialization at the myosin VI-actin interface and a mechanism of force sensitivity. eLife, 6:e31125, 2017.

[37] M Doi and SF Edwards. The Theory of Polymer Dynamics. Clarendon Press, Oxford, 2013.

